# Temporal and spatial modulation of the immune response of the murine Gl261 glioma tumor microenvironment

**DOI:** 10.1101/858894

**Authors:** Kelly J McKelvey, Amanda L Hudson, Ramyashree Prasanna Kumar, James S Wilmott, Grace H Attrill, Georgina V Long, Richard A Scolyer, Stephen J Clarke, Helen R Wheeler, Connie I Diakos, Viive M Howell

**Affiliations:** Bill Walsh Translational Cancer Research Laboratory, Kolling Institute, The University of Sydney Northern Clinical School and Northern Sydney Local Health District, St Leonards, NSW, Australia; Sydney Vital Translational Research Centre, Royal North Shore Hospital, St Leonards, NSW, Australia; The Brain Cancer Group, St Leonards, NSW, Australia; Melanoma Institute Australia, North Sydney, NSW, Australia; Charles Perkins Centre, The University of Sydney, Camperdown, NSW, Australia; Northern Sydney Cancer Centre, Royal North Shore Hospital, St Leonards, NSW, Australia; Mater Hospital, North Sydney, NSW, Australia; Royal Prince Alfred Hospital, Camperdown, NSW, Australia

## Abstract

Glioblastoma, the most aggressive form of glioma, has a 5-year survival rate of <5%. While radiation and immunotherapies are routinely studied in the murine Gl261 glioma model, little is known about its inherent immune response. This study quantifies the temporal and spatial localization of immune cell populations and mediators during glioma development.

Eight-week old male C57Bl/6 mice were orthotopically inoculated with 1×10^6^ Gl261 cells and tumor morphology, local and systemic immune cell populations, and plasma cytokines/chemokines assessed at Day-0, 1, 3, 7, 14, and 21 post-inoculation by magnetic resonance imaging, chromogenic immunohistochemistry, multiplex immunofluorescent immunohistochemistry, flow cytometry and multiplex immunoassay respectively.

From Day-3 tumors were distinguishable with >30% Ki67 and increased tissue vascularization (p<0.05). Increasing tumor proliferation/malignancy and vascularization were associated with significant temporal changes in immune cell populations within the tumor (p<0.05) and systemic compartments (p=0.02 to p<0.0001). Of note, at Day-14 16/24 plasma cytokine/chemokines levels decreased coinciding with an increase in tumor cytotoxic T cells, natural killer and natural killer/T cells. Data derived provide baseline characterization of the local and systemic immune response during glioma development. They reveal that type II macrophages and myeloid-derived suppressor cells are more prevalent in tumors than regulatory T cells, highlighting these cell types for further therapeutic exploration.

## 1 Introduction

High grade gliomas (HGG) remain a debilitating and early fatal disease. Almost 60% of gliomas are glioblastomas (GBM) which develop *de novo* (primary GBM), though secondary GBM can develop from the progression of lower grade gliomas (1). In Australia, glioblastomas account for ∼1,000 HGG cases per year and have a dismal prognosis of 12-16 months. While our understanding of the tumor biology has increased, patient outcomes have not improved significantly in the last 2 decades. At present the standard therapy for GBM remains maximum safe surgical resection, concomitant chemo-radiation, adjuvant chemotherapy, with/without the addition of anti-angiogenic drug bevacizumab (Avastin™) as a second line treatment. While this multi-modal therapy has brought a modest overall survival benefit in patients with temozolomide (TMZ)-treatable methylated O^6^-methylguanine-DNA methyltransferase (MGMT) promoter, it confers little benefit in the 50% of GBM patients with an unmethylated MGMT promoter (2). Greater therapeutic options are desperately required.

Immunotherapies have been suggested as a potential therapy in the notoriously immunogenically ‘cold’ GBM tumors. Vaccines (3, 4), oncolytic viruses (5) and cannabis compounds (THC and CBD; clinical trial NCT01812603) are showing promise in Phase II/III clinical trials. Yet, while our understanding of brain cancer tumor biology has advanced, the modulation of the anti-tumor immune response with these therapeutic interventions as single-modal agents or in combination with standard care remains comparatively poorly understood. As we move toward personalized medicine, our ability to monitor and track and individual’s immune response to the developing glioma and response to different therapeutic intervention will become of paramount importance to determine what therapy should be administered and when in order to support tumor stabilization/elimination. This has precedence. The IMMO-GLIO-01 glioma clinical trial published a case study of a 53-year old women with GBM in which they longitudinally monitored her immune response during the treatment regimen, noting that a shift of the CD4:CD8 ratio was associated with magnetic resonance imaging (MRI) tumor progression (6).

Intra-tumoral heterogeneity is a hallmark of GBM (7), and recurrence is associated with polarization toward an immunosuppressive microenvironment (8). Inflammation and coagulation play significant roles in tumor elimination, equilibrium and escape (9, 10) and by better understanding how standard therapy modulates this microenvironment (10, 11) we may better harness the immune system to treat these tumors. The use of animal models has transformed this area with the development of grafted syngeneic, immunocompromised and humanized patient-derived xenograft, and spontaneous genetically-modified or chemically-induced mouse models each with their own advantages and disadvantages (12-15). For example, they have revealed that genetic driver mutations create unique microenvironments in GBM (16, 17) and have enabled longitudinal and multiregional investigation of novel effective therapeutics on the tumor (18), where repeat needle biopsies in this patient population is fraught with challenges. In the interest of assessing immunotherapies and targeted therapies in brain cancer such models offer a valuable preclinical resource (19).

The murine Gl261 glioma model has been well utilized for novel therapeutics as a surrogate for brain cancer, yet remains poorly characterized from the tumor microenvironment and systemic immune response perspective. The current study builds upon the historical tumor biology reports in this model (20-24). The overall aim of this study was to characterize the spatial and temporal immune response and tumor microenvironment in the murine Gl261 glioma model, and determine whether plasma mediators correlate with tumor immunity. The findings confirm the dynamic processes of immune response during tumor development in both local and systemic compartments and show that at day 14 post-inoculation the decrease in plasma cytokines/chemokines correlates with an increase in cytotoxic T cells (Tc), natural killer (NK) and NK/T cells in the tumor infiltrate.

## 2 Materials and Methods

### 2.1 Mice

Murine glioma Gl261 cells were kindly donated by Géza Sáfrány (Frederic Joliot-Curie National Research Institute for Radiobiology and Radiohygiene, Hungary). The animal study was reviewed, approved and performed in accordance with the Northern Sydney Local Heath District Animal Ethics Committee guidelines, Royal North Shore Hospital, St Leonards, Australia (Approval #RESP/17/205). Eight week old male C57Bl/6.Kearn’s mice were stereotactically inoculated with 1×10^6^/2μl murine glioma Gl261 cells into the right caudoputamen (striatum) at mediolateral 2mm, anteroposterior −0.1mm, dorsoventral 2.6mm Bregma. Mice were kept on 12 hour day / night light cycles with standard chow and water *ad libitum*. Mice were randomly assigned into groups with 15 mice per group and monitored for well-being and euthanized by cardiac puncture at the pre-determined endpoints of 0, 1, 3, 7, 14, and 21 days after inoculation. *Ex vivo* magnetic resonance imaging (MRI) was performed on 3 mice per group, and histopathology and immune cell and cytokine analysis performed on 12 mice in each group.

### 2.2 MRI

Brains were harvested and fixed in 10% v/v neutral buffered formalin (NBF) for 24 hr and washed in phosphate buffered saline (PBS). *Ex vivo* single non-contrast T2-weighted 2D RARE quick scans were acquired using a Biospec Avance III 94/21 USR preclinical MRI (Bruker BioSpin) performed by the Biological Resources Imaging Laboratory, University of New South Wales. Echo time = 45 ms; repetition time = 2559.348 s; number of averages = 16; matrix = 256 × 128; 23 slices; slice thickness = 500 μm).

### 2.3 Histopathology

Tumor morphology and necrosis was assessed by Mayer’s hemotoxylin and eosin Y/erythrosin B staining.

Brains were harvested and fixed in 10% v/v NBF for 24 hr. Following paraffin embedding, 4μm sections were rehydrated and microwave antigen retrieval performed in citrate buffer, pH 6.0. Next, sections were blocked with 2.5% v/v normal goat serum, and incubated with primary antibody for 1hr at room temperature; CD31 (0.013μg/ml; 77699) and Ki67 (0.0835μg/ml; 12202; Cell Signaling Technologies). For detection, sections were incubated with ImmPRESS™ horse radish peroxidase (HRP) goat anti-rabbit immunoglobulin G (IgG) polymer (MP-7451; Vector Labs) for 30min at room temperature and visualized using NovaRed (SK-48000; VectorLabs).

Stained slides were digitally scanned using the Aperio AT2 Digital Pathology Scanner. For histopathological assessment five random digital images of the tumor region per slide was captured at 20x magnification using Aperio ImageScope (v12.3.2.8013; Leica Biosytems). Areas of normal brain tissue and tumor necrosis were excluded and then Ki67 positive staining was quantified by ImmunoRatio ImageJ plugin (v1.0c, 14.2.2011; http://jvsmicroscope.uta.fi/immunoratio/) and CD31 positive vessels enumerated and measured using the Microvessel-Segmentation MATLAB plugin (25).

### 2.4 Multiplex fluorescent immunohistochemistry

Sections were sequentially stained using the OPAL polymer anti-rabbit HRP (PerkinElmer) or ImmPRESS™ HRP goat anti-rat IgG polymer (MP-7404; Vector Labs) with the following primary antibodies for 1 hour at room temperature; Panel 1: CD3 (0.3μg/ml; A045201; DAKO), CD4 (0.31μg/ml; ab183685; Abcam), CD8 (0.50μg/ml; ab203035; Abcam), FoxP3 (0.05μg/ml; 12653S; Cell Signaling Technologies), CD161 (0.031μg/ml; ab234107; Abcam), and Ki67 (0.084μg/ml; 12202S; Cell Signaling Technologies); Panel 2: ARG1 (0.125μg/ml; ab91279; Abcam), iNOS (8μg/ml; ab3523; Abcam), F4/80 (0.22μg/ml; 70076S; Abcam), TMEM119 (0.076μg/ml; ab209064; Abcam), CD19 (0.02μg/ml; 90176S; Cell Signaling Technologies), and Ki67 (0.084μg/ml; 12202S;Cell Signaling Technologies); Panel 3: Ly6C (0.10μg/ml; ab15627), Ly6G (0.25μg/ml; ab25377; Abcam), CD11b (0.085μg/ml; ab133357; Abcam), F4/80 (0.22μg/ml; 70076S; Cell Signaling Technologies), GFAP (1.45μg/ml; Z033401-2; DAKO) and Ki67 (0.084μg/ml; 12202S;Cell Signaling Technologies). Six color OPAL reagents were utilized (1:100 OPAL 520, 540, 570, 620, 650, 690) and all sections were counterstained with DAPI for nuclei. Sections were scanned at 10x, regions of interest denoted in Phenochart version 1.0.2, and then multi-spectral images (MSI) acquired at 20x using a VECTRA 3.0 (PerkinElmer). Spectral unmixing and cell phenotypes were quantified using InForm version 2.4 (PerkinElmer) with technical assistance from the Sydney Microscopy and Microanalysis Unit, University of Sydney.

Immune cell populations were defined as CD3^+^ T cell, CD3^+^CD4^+^ helper T-lymphocytes (Th), CD4^+^FOXP3^+^ regulatory T cells (Treg), CD3^+^CD8^+^ Tc, CD161^+^ NK cells, CD3^+^CD161^+^ NK/T, F4/80^+^ macrophage (Mac), F4/80^+^iNOS^+^ type I macrophage (M1), F4/80^+^ARG1^+^ M2, TMEM119^+^ microglia, CD11b^+^ dendritic cell (DC), CD11b^+^Ly6C^high^ monocytic-myeloid derived suppessor cells (M-MDSC), CD11b^+^Ly6G^high^ polymorphonuclear (PMN)-MDSC, and CD19^+^ B cells expressed as mean number of cells per high power field (HPF).

### 2.5 Hematology and Flow cytometry

Whole blood (1ml) was collected via cardiac puncture into K_3_EDTA tubes (Minicollect®; Greiner Bio-One) and red blood cell, lymphocyte and platelet counts acquired by COULTER® Ac-T diff hematology analyzer with the Vet App 1.06 (Beckman Coulter).

To assess systemic immune cell populations multi-colour flow cytometry was employed. Using 100μl whole blood, 1×10^6^ splenocytes and 1×10^6^ bone marrow-derived cells, red blood cell were lysed and leukocytes stained with a cocktail of antibodies; CD25-BV421 (1μl; 564370), FV510-BV510 (1μl; 564406), CD80-BV605 (1μl; 563052), NK1.1-BV650 (1μl; 564143), CD4-BV711 (0.25μl; 563726), CD117-BV786 (1μl; 564012), CD11b-BB515 (0.25μl; 564454), CD19-PerCP/Cy5.5 (1μl; 551001), CD115-PE (0.25μl; 565249), Ly6G-PE/CF594 (0.06μl; 562700), CD3-PE/Cy7 (1μl; 552774), CD206-AF647 (1μl; 565250), CD8a-AF700 (0.25μl; 557959), Ly6C-APC/Cy7 (0.5μl; 560596; all BD Biosciences). Data was acquired using a BD LSRFortessa™ and analyzed using BD FACSDiva™ Software version 6 (BD Biosciences).

Immune cell populations were defined as CD3^+^ T cell, CD3^+^CD4^+^ Th, CD3^+^CD4^+^CD25^+^ Treg, CD3^+^CD8 Tc, CD3^-^NK1.1^+^ NK, CD3^+^NK1.1^+^ NK/T, CD115^+^CD11b^+^ monocytes, CD115^+^CD11b^+^CD80^+^ M1, CD115^+^CD11b^+^CD206^+^ M2, CD115^-^CD11b^+^ DC, CD115^-^ CD11b^+^Ly6C^high^Ly6G^-^ M-MDSC, CD115^-^CD11b^+^Ly6C^low^Ly6G^high^ PMN-MDSC, CD117^+^hematopoietic stem cell (HSC), CD19^+^ B cells and expressed as a percentage of the parent population.

### 2.6 Multiplex immunoassays

Plasma was obtained from whole blood by centrifugation at 500 x g for 5 minutes at room temperature. Mouse cytokine 23-plex immunoassay (Bio-Plex®; Bio-Rad Laboratories) and chromogenic sandwich enzyme-linked immunosorbent assay (ELISA) for TGF-β1 (DY1679; R&D Systems) were performed in accordance with the manufacturer’s instructions.

### 2.7 Statistical analyses

Histological data are expressed as the mean of 5 high power fields ± standard error of the mean (SEM) and assessed for significance using One Way Analysis of Variance (ANOVA) with Tukey’s Multiple Comparison Test. Hematology, flow cytometry and chemokine/cytokine data are expressed as median ± interquartile range due to significant differences in standard deviation for some parameters, as determined by the Brown-Forsythe Test for normality. Kruskal-Wallis One Way ANOVA with Dunn’s Multiple Comparison Test was performed using Prism 7 for Windows (GraphPad Software, Inc). Multiplex immunofluorescent data are expressed as the mean of all 20x high power MSI fields covering the tumor region (or equivalent brain region in D0 sham controls) ± SEM. Statistical significance was assessed by One Way ANOVA with Tukey’s Multiple Comparison Test. Relationships between plasma cytokine levels and tumor infiltrate were assessed by Pearson correlation. All statistical analyses were performed using Prism 7 for Windows (GraphPad Software, Inc) and statistical significance deemed p-value less than 0.05.

## 3 Results

### 3.1 MRI demonstrated visible tumor growth from day 14 with mass effect and contralateral oedema

While the needle tract was evident at the early time points, definitive tumor morphology was only distinguishable on the non-contrast T2-weight scans from day 14 post-inoculation (Fig 1). Tumors grew posteriorly into the hippocampal and mid brain regions and were associated with mass effect and a midline shift at day 14 (Fig 1). Marked brain edema in the contralateral hemispheres of brains from the day 21 cohort was also observed. Dark (hematoma) and brighter intensity (fluid-filled) regions were also observed in the central region of the tumors at day 21 post-inoculation.

**Fig 1.**
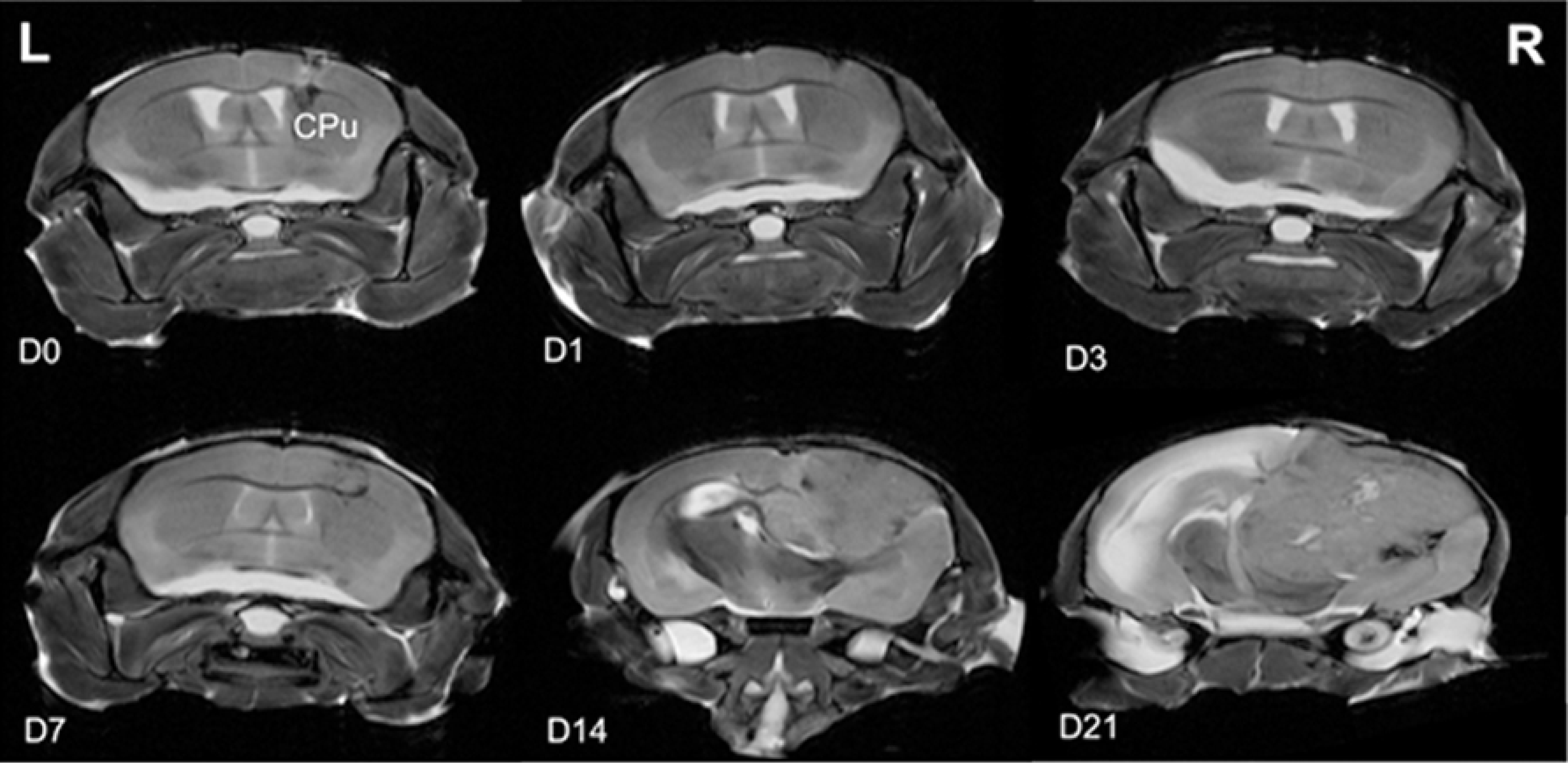
Ex vivo non-contrast T2-weighted 2D MRI scans of ex vivo brains depicting the tumor progression at day 0 (D0), D1, D3, D7, D14, and D21 post-inoculation of murine glioma Gl261 cells into the caudoputamen (CPu). Left (L) and right (R) of the brain are indicated. Images are representative of N = 3 mice per time point.

### 3.2 Tumors showed increasing proliferative rates, blood vasculature and geographic necrosis

Hematoxylin and eosin-based tissue morphology demonstrated the progressive infiltration and development of the glioma within the caudoputamen region (Fig 2A). Sham-operated (D0) brains showed a region of hemorrhage at the margin of the needle tract, which largely resolved by day 1-3. At day 3 the cells show uniform shape and nuclei, which progressed to an elongated, morphology at days 7 and 14. At day 21 the cells are densely packed and show heterogeneous cellular size and nuclear formations (Fig 2B).

**Fig 2.**
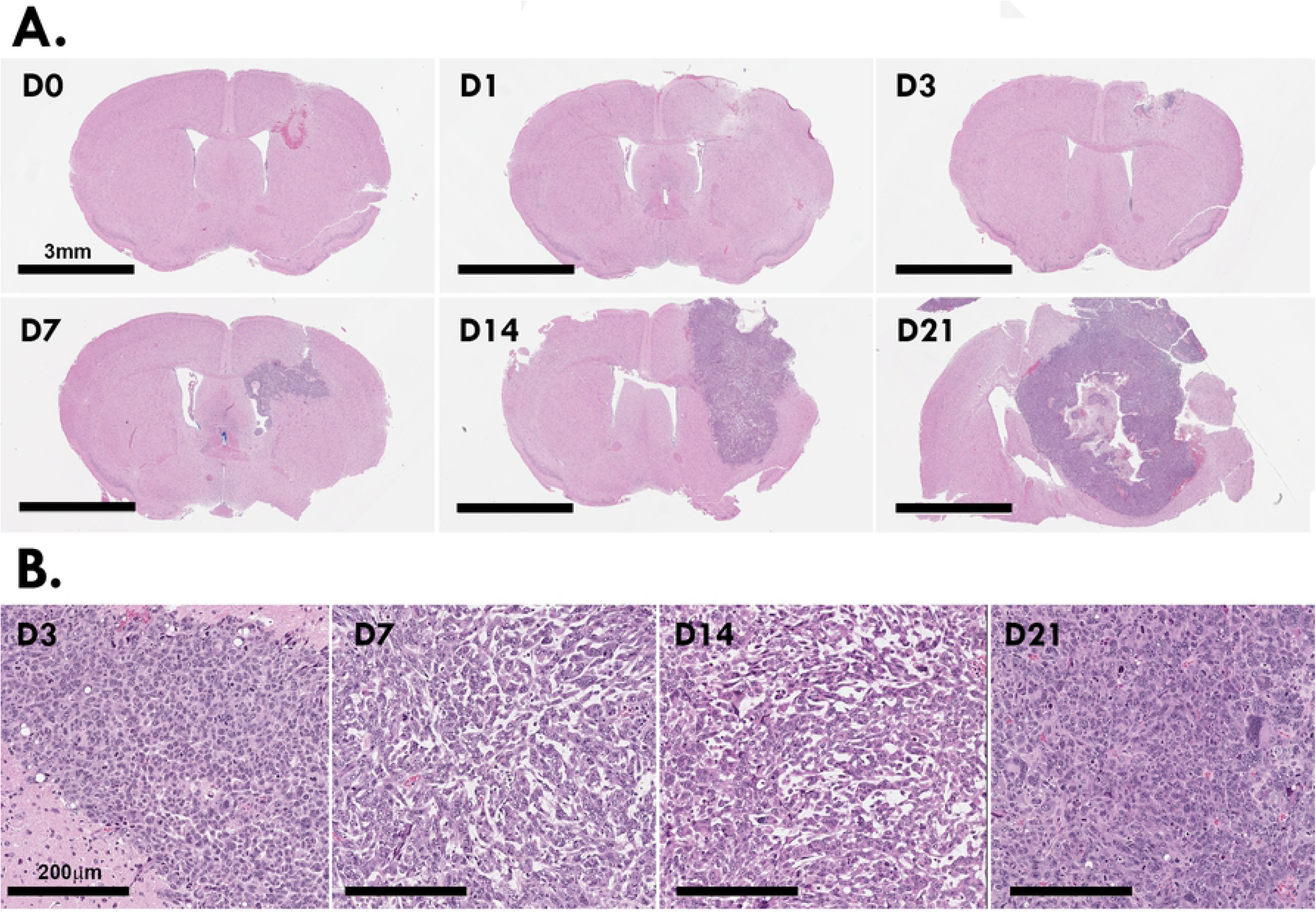
4μm paraffin sections of murine Gl261 brain tumors stained with hematoxylin and eosin Y/eythrosin (**A**). High resolution of morphology of the Gl261 cells as the tumor grows and matures (**B**). Images are representative of N = 12 mice per time point.

Quantification of Ki67 staining showed increasing cell proliferation within the tumor microenvironment up to day 14 (Fig 3A). As the tumors enlarged beyond day 14, centralized regions showed evidence of necrosis. At day 21, where a centralized foci of geographic necrosis was present, Ki67 positivity of the surrounding tumor tissue decreased (Fig 3B). Cell proliferation peaked from day 3-14 post-inoculation (mean ± SEM; D3 32.99 ± 3.68%; D7 37.78 ± 3.74%; D14 37.95 ± 2.09%) compared to sham-operated (D0 4.58 ± 0.50%; Fig 3C). The ∼2-fold decrease in Ki67 compared to the peak and the presence of necrotic regions at day 21 indicate that the tumor has exceeded its blood supply.

**Fig 3.**
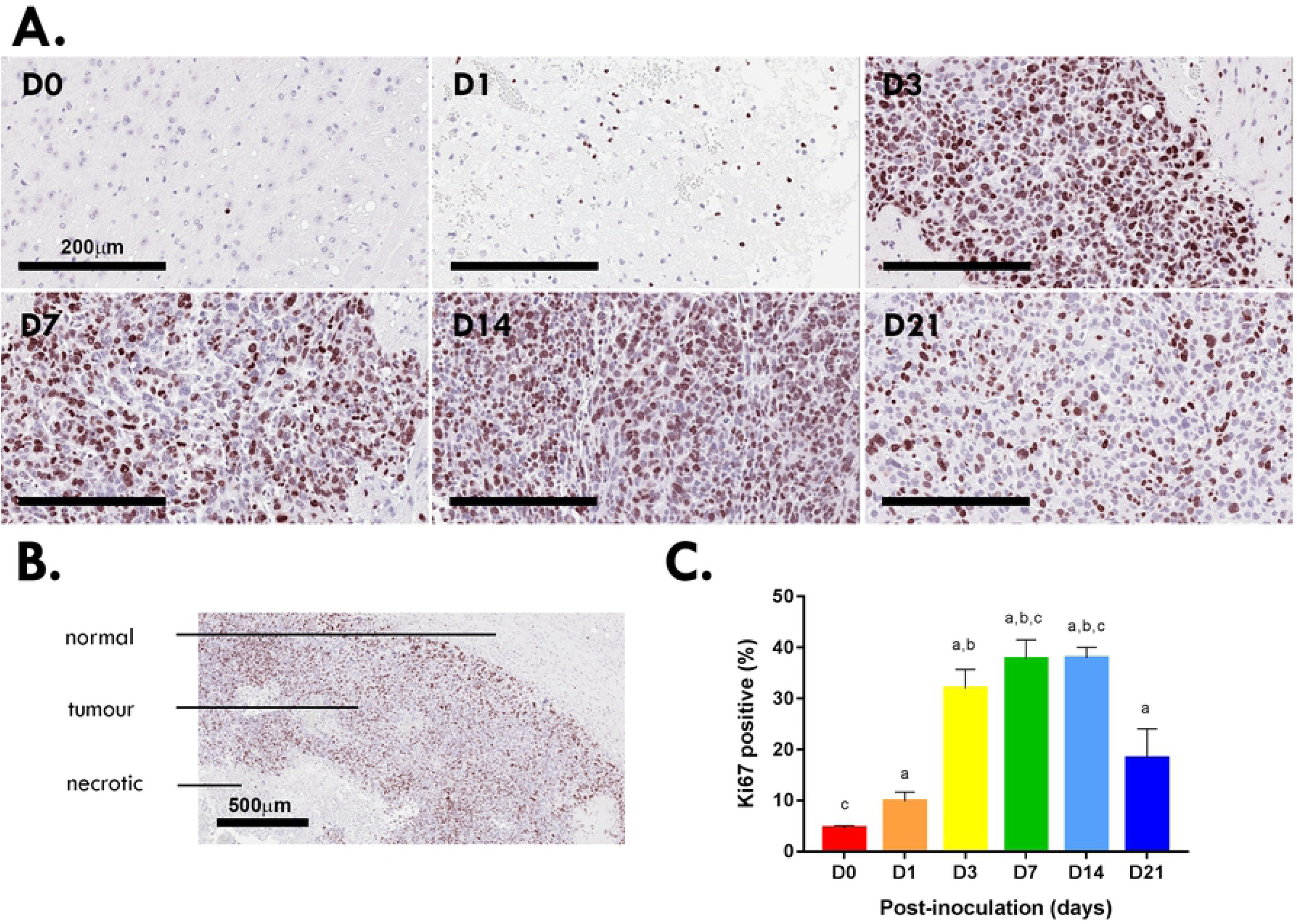
Ki67^+^ staining at the inoculation site of murine Gl261 gliomas (**A**). Scale bar, 200μm. Tumors at day 21 post-inoculation showed signs of hypoxia/necrosis (**B**). Scale bar, 500μm. Quantitation of five high power fields for each murine brain (N=12 per time point; **C**). Data are expressed as mean ± SEM per high power field. ^a^p<0.05 vs D0, ^b^p<0.05 vs D1, ^c^p<0.05 vs D21 by Tukey’s multiple comparison test.

To confirm that blood vasculature within the tumor contributed to the necrotic regions, the number and size (lumen area) of CD31^+^ blood vessels was quantified within the tumor microenvironment (Fig 4A). The number of vessel peaked at day 3 (26.38 ± 1.45) and then significantly declined at day 7 (17.11 ± 3.05) and 14 (13.77 ± 1.99; Fig 4B). At day 21 a significant increase in blood vessels and lumen area was present (Fig 4C). Other vasculature parameters assessed were vessel thickness, perimeter and area - but were not significantly different between time points.

**Fig 4.**
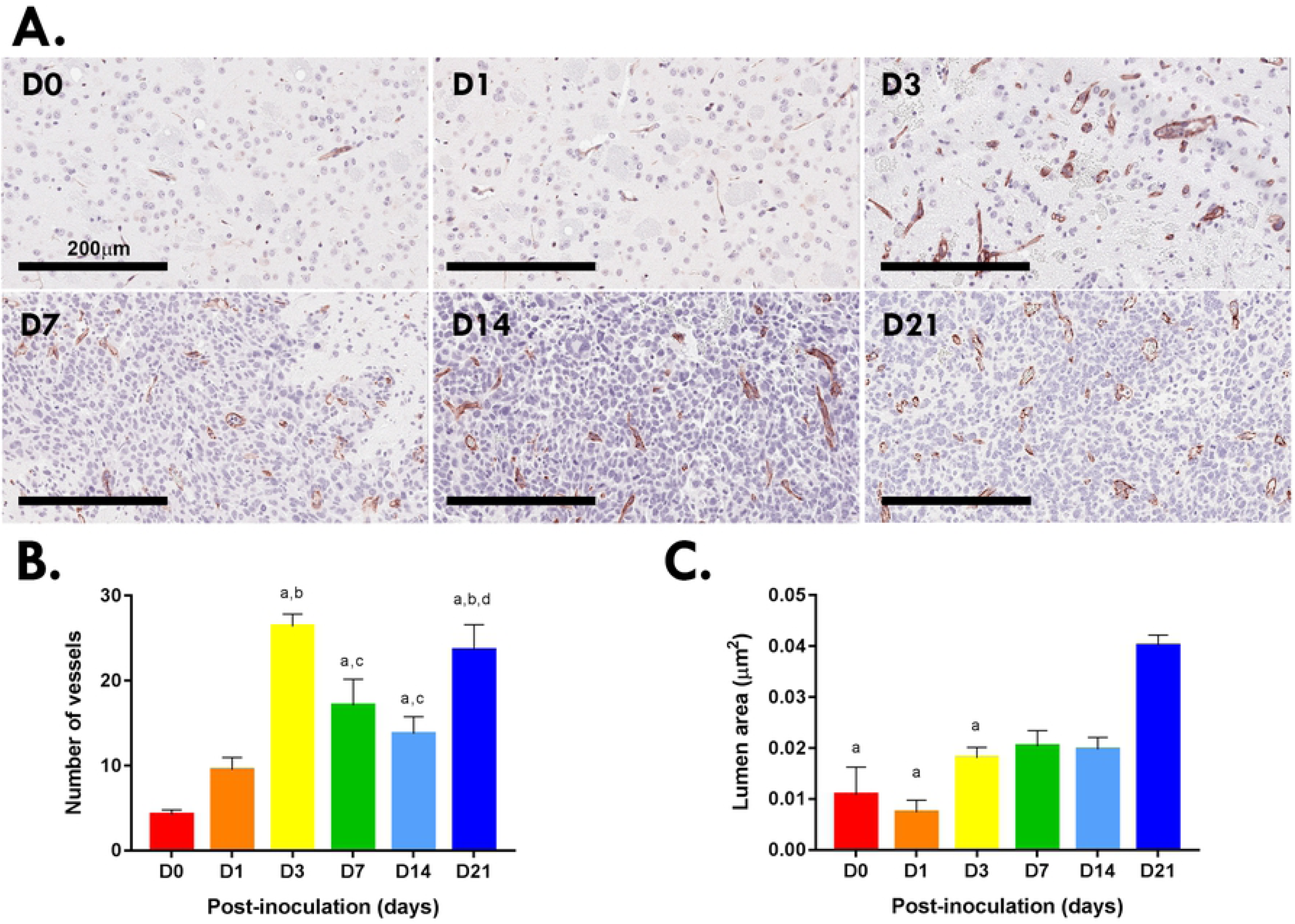
CD31^+^blood vessels at the inoculation site of murine Gl261 gliomas (**A**). Scale bar, 200μm. Quantitation of blood vessels (**B**) and lumen area (**C**) in five high power fields (20x) for each murine brain (N=12 brains per time point; **B**). Data are expressed as mean ± SEM per high power field. (**B**) ^a^p<0.05 vs D0, ^b^p<0.05 vs D1, ^c^p<0.05 vs D3, ^d^p<0.05 vs D14 by Tukey’s multiple comparison test. (**C**) ^a^p<0.05 vs 21 by Tukey’s multiple comparison test.

### 3.3 M2 macrophages and MDSCs are more prominent than Tregs in the tumor immune infiltrate

Three multiplex antibody panels were used to assess the infiltration of lymphocytes (Panel 1), macrophages (Panel 2) and DC (Panel 3) populations into the brain tumors using multiplex fluorescent immunohistochemistry (Fig 5). Quantitative assessment showed that T and B cell populations increased significantly over time, with Tregs peaking at day 7 (mean ± SEM; 0.20 ± 0.00 cells per HPF) compared to days 0 and 21 where Tregs were absent (both 0.00 ± 0.00 cells per HPF; p<0.05). CD4^+^ Th, CD8^+^ Tc, NK and NK/T cells showed no significant differences over time. Mac, M2, and microglia increased over time peaking at day 14, while M1 spiked at days 7 (2.72 ± 0.02 cells per HPF; p<0.0001) and 21 (4.20 ± 0.01 cells per HPF; p<0.0001) compared to day 0 (0.06 ± 0.00 cells per HPF). Among the DC populations, DC, M-MDSC and PMN-MDSC populations significantly increased from day 3 and spiked at day 21 (DC 18.62 ± 0.07, M-MDSC 7.88 ± 0.03, PMN-MDSC 11.44 ± 0.04 cells per HPF; all p<0.0001). Surprisingly immunosuppressive populations did not all appear/disappear at the same time or to similar levels; Tregs peaked at day 7 but remained at markedly lower levels than M2 macrophages and MDSCs which peaked at day 14 and day 21 respectively.

**Fig 5.**
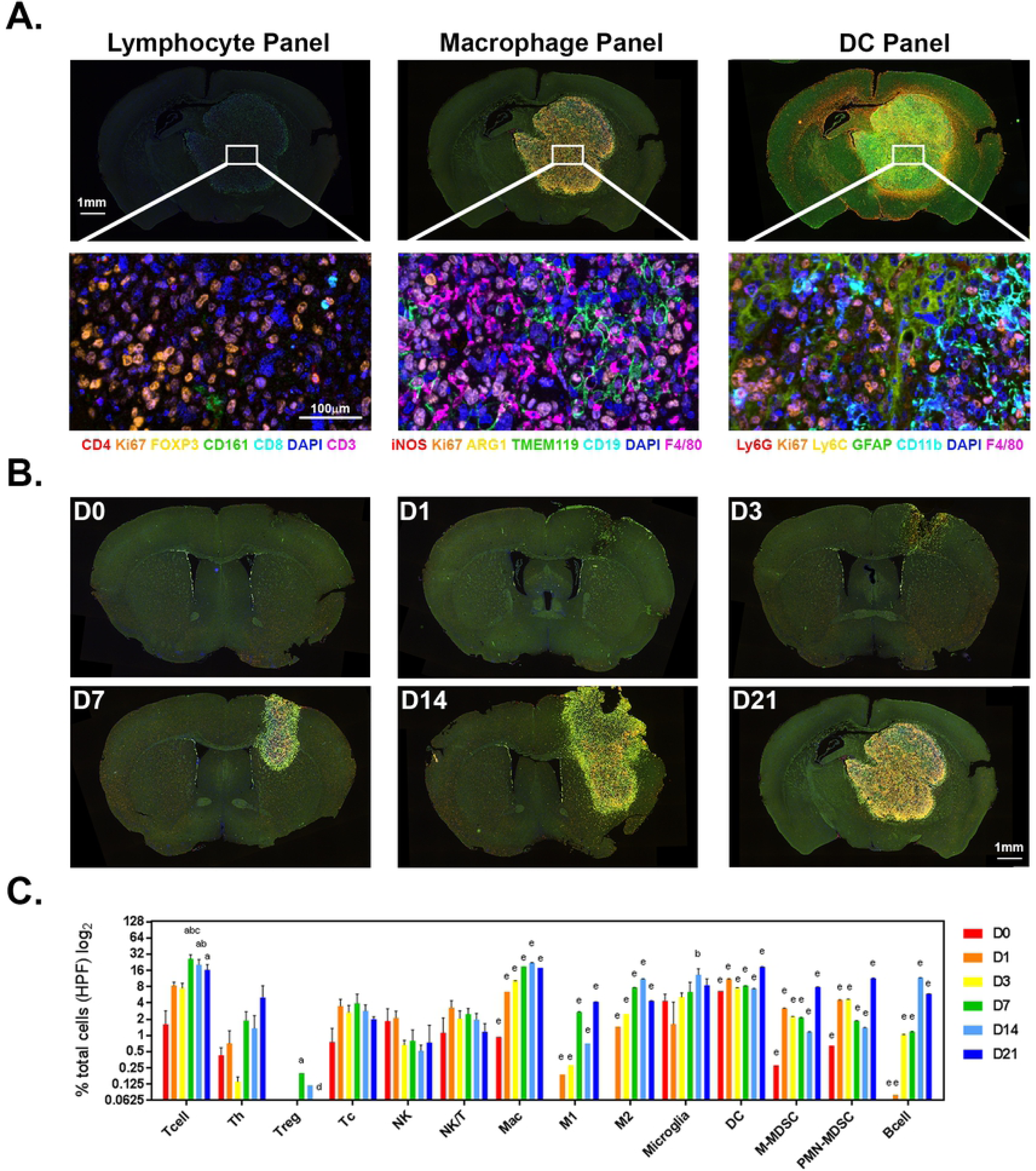
(**A**) Whole slide scans (4x) and MSI spectrally unmixed and pseudo-colored micrographs (20x) of the three immune cell panels for lymphocytes, macrophages and DC populations in the tumor infiltrate. (**B**) Whole slide scans of multiplex immunofluorescent stained murine brain murine Gl261 gliomas over time depicting the macrophage antibody panel. (**C**) Quantitation of immune cell phenotypes in high power fields for each murine brain (N=6 brains per time point). Data are expressed as mean ± SEM per high power field. ^a^p<0.05 vs D0, ^b^p<0.05 vs D1, ^c^p<0.05 vs D3, ^d^p<0.05 vs D7, ^e^p<0.05 vs all other time points by One-way ANOVA with Tukey’s multiple comparison test.

### 3.4 Systemic immunity to glioma development is dynamic but does not directly correlate with the tumor infiltrate

To assess the systemic immune response, hematological parameters and immune cell populations in the spleen, bone marrow and peripheral blood were quantified by flow cytometric analysis. In response to the tumor inoculation wound, platelets decreased by 20% at day 1 compared to day 0 (p<0.005) but normalized by day 3 (Fig 6A). Total circulating white blood cells were increased at day 14 (Fig 6A) and further delineated for temporal modulation of the immune cell populations in peripheral blood, as well as the spleen and bone marrow compartments. The gating strategy for delineation of the immune cell populations is shown in Fig 6B.

**Fig 6.**
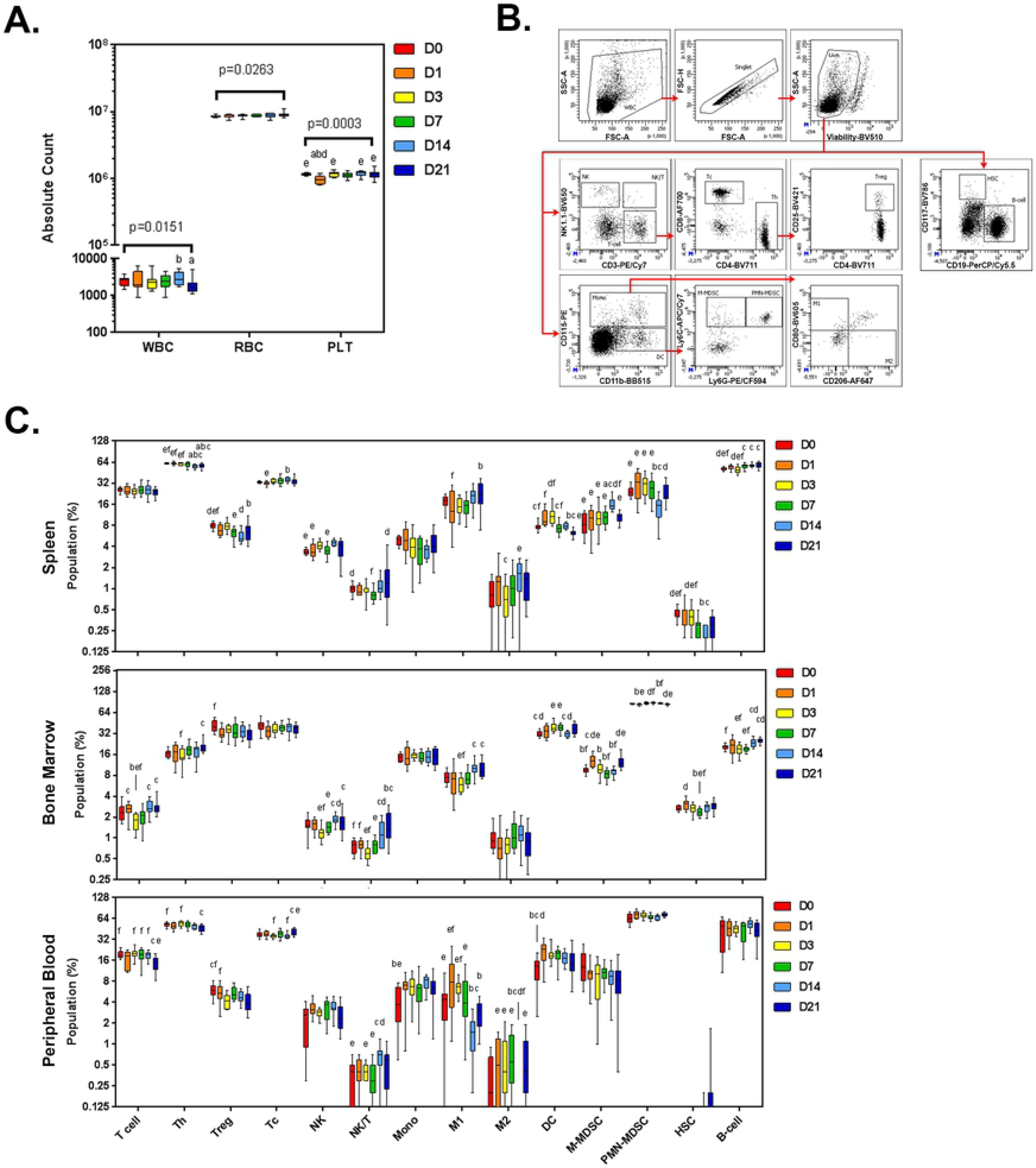
Systemic immune cell populations. Hematology (**A**) and immune cell populations from spleen, bone marrow and peripheral blood from mice inoculated with glioma Gl261 cells. Cells were harvested at day 0 (sham) to day 21 time points and assessed by multi-color flow cytometry. Gating strategy is depicted in **B**, and quantitative bar graphs in **C**. Data are expressed as absolute count (**A**) and percentage of parent population (%; **C**). ^a^p<0.05 vs D0, ^b^p<0.05 vs D1, ^c^p<0.05 vs D3, ^d^p<0.05 vs D7; ^e^p<0.05 vs D14; ^f^p<0.05 vs D21 by Dunn’s Multiple Comparison Test.

Within the spleen, Tregs and PMN-MDSC significantly decreased by ∼10% and ∼40%, respectively, at day 14 post-inoculation, while Tc, M2 and M-MDSCs all peaked at day 14 when compared to other time points assessed (Fig 6C).

In patients with glioma, immune cell populations are reported to sequester in the bone marrow compartment (26). Consistent with this notion, the data here show that in bone marrow, ‘anti-tumor’ populations T cell, Th, NK, NK/T, M1 and B cells all decreased at day 3 and highest at day 21 (Fig 6B) potentially suggesting progressive sequestration within the bone marrow compartment, though local expansion of these immune cells populations is also plausible. In the immunosuppressive populations, Tregs decreased at day 21 compared to day 0, meanwhile the MDSC populations showed an inverse relationship to one another; M-MDSCs were lowest at day 7 and peaked at day 1 and 21, yet PMN-MDSCs peaked at day 7, and were lowest at day 1 and 21 (Fig 6B).

In peripheral blood, Th and Treg populations decreased 12% and 30% at day 21 compared to day 0, respectively (Fig 6B). NK/T peaked at day 14 and Tcs at day 21. A perplexing result was the significant increase in monocytes at day 14, yet M1 cells had declined and M2 were entirely absent (Fig 6B). Correlation of the systemic peripheral blood immune cell populations quantified by flow cytometry, and immune cell infiltrate assessed by multiplex fluorescent immunohistochemical staining showed no significant correlations, except for M1 macrophages which showed a moderate correlation (r^2^=0.3849, p=0.0003; Pearson’s correlation). This suggests that non-invasive assessment of peripheral blood immune cell populations does not reflect the tumor infiltrate in this model.

### 3.5 Plasma cytokines, chemokines and growth factors correlate with an increase in Tc, NK and NK/T cells at day 14

To determine whether the change in immune cell response is indicated by a change in plasma cytokine levels, a panel of 24 cytokines was assessed by multiplex immunoassay or ELISA. The results are shown in Fig 7. At day 1, G-CSF increased 4.5-fold compared to day 0 consistent with a role in neutrophil mobilization and an early wound healing response. The pleiotropic cytokine, IL-6 was also increased ∼3-fold at day 1 compared to sham-operated animals (day 0). At day 14, 16 of the 24 cytokines, chemokines and growth factors assessed were significantly decreased; IL-1β, IL-3, IL-4, IL-6, IL-9, IL-10, IL-12(p40), IL-12(p70), IL-13, IFN-γ, TNF-α, G-CSF, GM-CSF, CCL2 and CCL3 (Fig 7).

**Fig 7.**
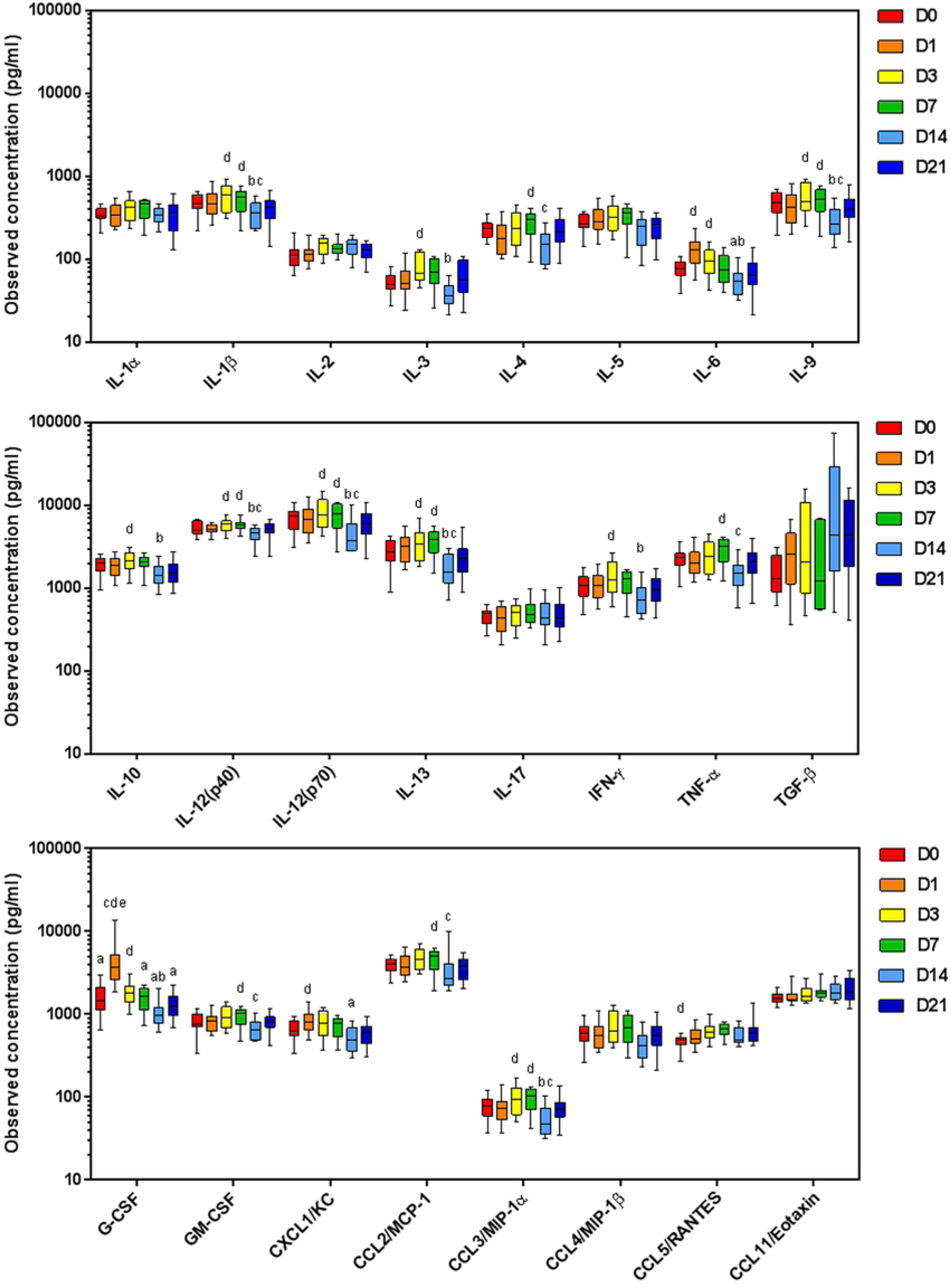
Plasma cytokine and chemokines levels from mice inoculated with glioma Gl261 cells. Plasma was harvested at day 0 (sham) to day 21 time points and assessed by 23-plex immunoassay, except TGF-β1 (by chromogenic ELISA). Data are expressed as observed concentration in pg/ml. ^a^p<0.05 vs D14; ^b^p<0.05 vs D21; ^c^p<0.05 vs D7; ^d^p<0.05 vs D3; ^e^p<0.05 vs D1 by Dunn’s Multiple Comparison Test.

Correlation studies of the plasma cytokines levels and tumor infiltrates for each animal revealed that as the above cytokines decreased at day 14, there was an increase in Tc, NK and NK/T cells in the tumor (r^2^ = 0.92 to 0.99; p= 0.001 to 0.04; Fig 8A). In contrast, CCL11 and TGF-β levels increased with an increase in these immune cell types in the immune infiltrate (r^2^ = 0.095 to 0.99; p= 0.001 to 0.03), as well as in NK and monocyte populations in the bone marrow (Fig 8B). Plasma IL-6 and G-CSF positively correlated with splenic DC and PMN-MDSC at all time points, M-MDSC were negatively correlated, consistent with roles in MDSC phenotypic development (Fig 8B).

**Fig 8.**
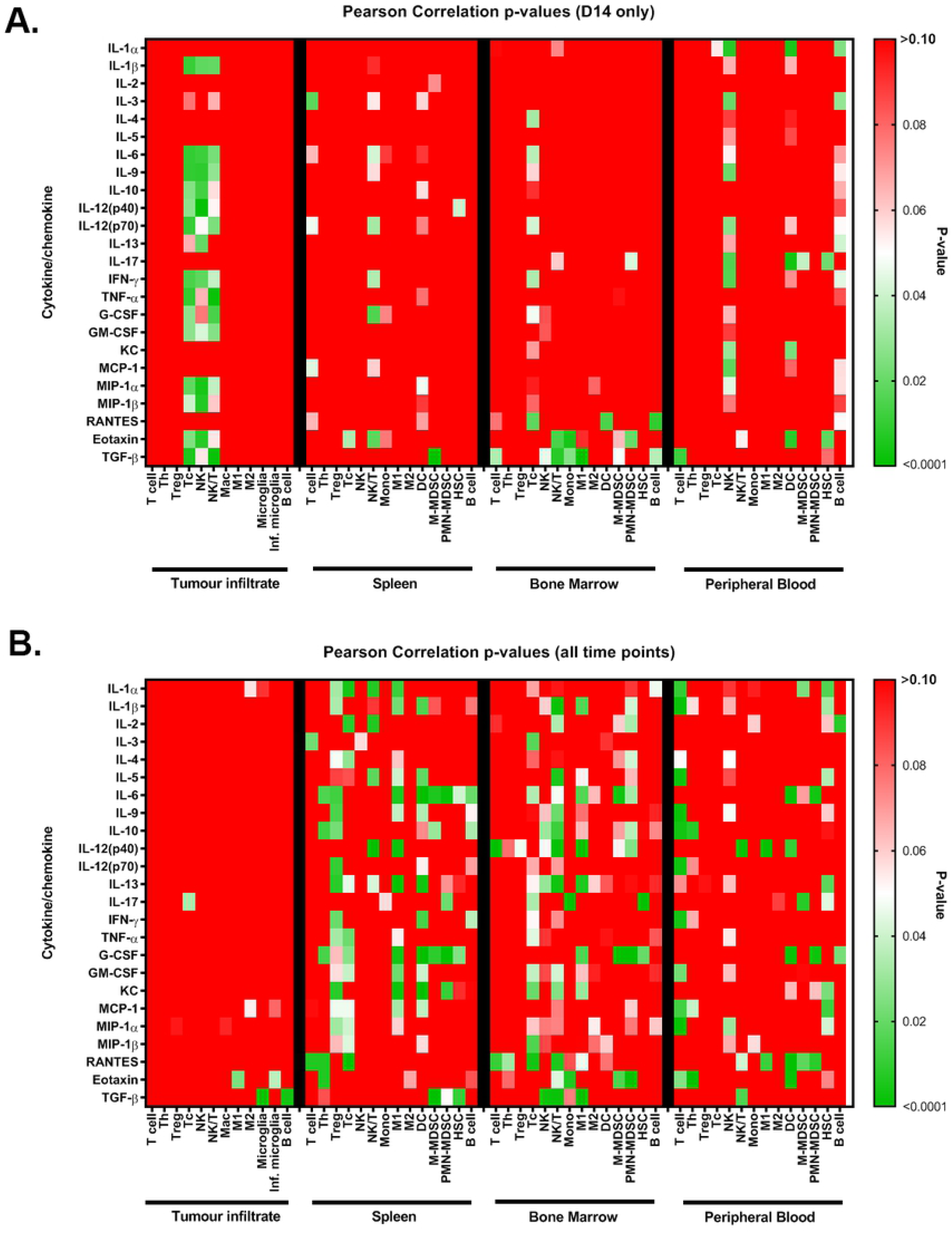
Colour maps of the p-values for Pearson correlations between plasma cytokine levels and tumor and systemic immune cell populations at day-14 post-inoculation (**A**) and at all time points (**B**). Red, p>0.05, white p=0.05, green p<0.05.

## 4 Discussion

GBM remains a debilitating and fatal cancer with a short median survival and limited standard treatment options. While a range of novel therapeutics are being examined, including but not limited to vaccines, oncoviruses, metabolism inhibitors, radiosensitizers/radioenhancers, nanoparticles, and immunotherapies, there remains a relative paucity of information on how the immune response responds to a developing glioma in the absence of therapeutic intervention. In this study we provide a comprehensive characterization of the immune response in the murine syngeneic Gl261 glioma model both temporally and spatially. This is the most widely used mouse model for GBM and with the increasing interest in immunotherapeutics, ‘point of reference’ knowledge of the inherent immune response within this model is central to the interpreting efficacy studies. Tracking the immune response to glioma and subsequent treatment therapies will additionally aid the design of optimal treatment strategies.

Using non-contrast T2-weighted MRI *ex vivo*, the Gl261 tumors demonstrated mass effect with midline shift and contralateral edema from day 14. While *ex vivo* MRI was used in this study as an observational tool to compare the clinical situation, current work in the area of preclinical medical imaging is combining MRI/positron-emission tomography (PET) tracers to solve the problem of the distinction of pseudo-response and -progression from tumor progression. In the orthotopic Gl261 model the combination of i.v. contrast MRI and [^18^F]-2-fluoro-d-(arabinofuranosyl) cytosine PET/CT enabled the distinction of regions of high immune inflammatory activity/infiltration in animals treated with DC vaccine and/or anti-programmed cell death-1 mAb (27), closing the gap between preclinical models and clinical treatment/monitoring.

Histopathologically the murine Gl261 tumors are poorly differentiated similar to human GBM cells (20, 21) with individual cell infiltration and invasion millimeters from the tumor boundary into normal brain tissue (23, 24). Consistent with other reports of Gl261 tumor morphology, from day 3 Gl261 cells showed uniform size and shape which progressed to elongated cells at day 7-14 with perivascular satellitosis, and increasing microvascular density and Ki67 proliferation indices (28, 29). At day 21 cells and nuclei showed heterogenic size and shape and the presence of centralized necrosis (23, 24, 28).

Despite Gl261 tumors being noted as partially immunogenic, expressing MHC I but low levels of MHC II, B7-1 and B7-2 (30), we saw no significant differences in ‘anti-tumor’ Th, Tc, NK and NK/T immune cell populations infiltrating the tumor microenvironment over time, although a correlation with decreasing plasma cytokines was noted at day 14 post-inoculation. TMEM119 was used to identify resident microglia from infiltrating macrophages in murine and human central nervous system tissue (31, 32), and showed little variation over time indicating that the resident population does not expand, but infiltrating macrophages contribute to the immune response. Among the ‘immunosuppressive’ populations, M-MDSC and PMN-MDSC significantly increased at day 3, then declined; Tregs spiked at day 7, and M2 at day 14. Finally, M-MDSC and PMN-MDSC spiked at day 21 when Tregs were absent. As aforementioned the marked decrease in 16 of 24 plasma cytokines and chemokines at day 14 correlated with an increase in pro-tumor cell types Tc, NK and NK/T cells, potentially indicating a re-activation of the immune response at this time point. We tentatively speculate that this may be associated with tumor immune escape (33), as the histological and MRI tumor growth appeared to accelerate around this time point.

The inter-relationship of Tregs, DCs and macrophages in glioma is complex. An absence of Tregs causes a phenotypic shift in macrophage populations toward M1 (34), while macrophages and microglia produce CCL2 within the glioma microenvironment, a chemokine that is critical for the recruitment of Treg and M-MDSCs (35). Gl261 tumors grown in CCL2-deficient mice or mice treated with a small-molecule antagonist of CCL-2 receptor, CCR4, failed to maximally accumulate Tregs and M-MDSCs within the glioma microenvironment (35). In our study plasma CCL2 levels peaked on day 7 and were markedly reduced at day 14 post-inoculation coinciding with changes in MDSCs within the tumor microenvironment, but not Tregs. Recent studies have shown bidirectional regulation of Treg cells and MDSCs directly through TGF-β (36), programmed cell death-1 (PD-1) and its ligand PD-L1 (37), and indirectly by controlling differentiation of Treg and regulatory DCs (38). Plasma TGF-β1 increased day 14 and 21 though was not significantly different due to heterogeneous levels, but correlated with tumor microglia across all times points.

In line with the upsurge in immunotherapy success in certain cancers, those which target T cell populations have become the focus of preclinical Gl261 studies of late. Checkpoint inhibitors PD-1, PDL-1 and TIM3 which regulate self-tolerance and Th1 responses respectively, and the rate-limiting metabolic enzyme and Treg modulator indoleamine 2,3-dioxygenase doubled median survival and increased long-term survival in cohorts when combined with acute radiation (39-41). The effect required continuous CD4^+^ T cell involvement (41). Interestingly the inclusion of TMZ chemotherapy with the trimodal anti-PD1, anti-TIM3 and radiation therapy reduced the number of long-term survival animals by 90% (41). The effect of TMZ on T cell populations is described elsewhere (42), though more data is needed in regards to combination therapy with novel immunotherapeutics.

In conclusion, the current study shows that there are dynamic immunomodulatory effects in the local and systemic compartments during glioma development and warrants comprehensive tracking during the preclinical assessment of novel therapeutics. Early data from the IMMO-GLIO-01 glioma clinical trial which tracks the immune response during treatment regimens demonstrates the value of such information. In a case study, RT treatment coincided with declines in T cells, B cells, pDCs and NK cells, and a shift of the CD4:CD8 ratio correlated with MRI tumor recurrence offering some hope in the identification of treatment response or prognostic indicators (6). In the move toward personalized medicine, tracking of the patient immune response before, during and after treatment and with different treatment combinations will give greater understanding of when and what therapy should be administered at each time point to induce/maintain a state of tumor stabilization/elimination. The current preclinical study aids this notion by providing critical insight into the dynamic immune response to glioma, the immunosuppressive immune cells that predominate the tumor infiltrate (M2 and MDSC vs Treg), and that changes in plasma cytokine profiles may indicate changes in the tumor immune infiltrate. Further exploration of the dynamic immune response in the context of novel therapeutics for glioblastoma is warranted in this model.

## 5 Acknowledgments

KM is supported by the Matt Callander ‘Beanie for Brain Cancer’ HMRI Fellowship funded by the Mark Hughes Foundation and AH by The Brain Cancer Group Fellowship. The project was supported by project grants from The Mark Hughes Foundation, The Brain Cancer Group, and Sydney Vital Translational Research Centre. Purchase of the VECTRA3 microscope was supported by The James N Kirby Foundation and The Goodridge Foundation.

We thank the following units for technical assistance; Biological Resources Imaging Laboratory, University of New South Wales (MRI); Histology Services, Hunter Medical Research Institute (histopathology); and Sydney Microscopy and Microanalysis Unit, The University of Sydney (multiplex fluorescent immunohistochemistry).

## 7 Author Contributions

KM: Conceptulization, Formal Analysis, Investigation, Methodology, Project Administration, Visualisation, Writing – Original Draft Preparation, Writing – Review & Editing

AH: Conceptulization, Investigation, Writing – Review & Editing

RPK: Investigation, Writing – Review & Editing

JW: Methodology, Resources, Writing – Review & Editing

GA: Methodology, Resources, Writing – Review & Editing

GL: Resources, Writing – Review & Editing

RS: Resources, Writing – Review & Editing

SC: Conceptulization, Supervision, Writing – Review & Editing

HW: Conceptulization, Supervision, Writing – Review & Editing

CD: Conceptulization, Funding Acquisition, Supervision, Writing – Review & Editing

VH: Conceptulization, Funding Acquisition, Supervision, Writing – Review & Editing

